# Patterns of genAI bias in guiding prospective undergraduate students: a study of UK neuroscience programmes

**DOI:** 10.64898/2026.03.20.713226

**Authors:** Harry G. Potter

## Abstract

Generative artificial intelligence (genAI) tools are increasingly used by prospective higher education (HE) applicants seeking guidance on university and programme selection. Despite rapidly expanding use, little is known about how genAI systems may introduce or amplify bias in undergraduate admissions decision-making. Here, we systematically examined patterns of bias across three widely used genAI chatbots (ChatGPT, Copilot, Gemini) using neuroscience as a representative UK undergraduate programme.

We constructed 216 prompts that varied by applicant characteristics (e.g. gender, study type, academic attainment). Each prompt was submitted to all three chatbots, generating 648 responses and 3240 individual programme recommendations. Output responses underwent text analysis (e.g. n-grams, gender-coded language), and national HE markers of esteem (REF21, TEF23, NSS24) were analysed.

Applicant grades and priorities produced the strongest effects on genAI outputs. Higher-grade applicants and those prioritising research received significantly more masculine-coded language, independent of applicant gender. N-gram patterns also diverged: high-grade prompts more frequently elicited terms relating to excellence and research intensity, whereas lower-grade prompts produced greater emphasis on widening access. Recommendations were systematically skewed, with higher grades, private schooling, and research-focused priorities increasing the likelihood of recommending elite institutions and programmes with higher entry requirements. Critically, the gender-coded language of outputs predicted institutional characteristics: masculine-coded responses were associated with recommendations featuring higher entry thresholds and stronger research performance, while feminine-coded responses favoured institutions with higher student satisfaction.

These findings reveal clear, systematic biases in how genAI guides prospective HE applicants. Such biases risk reinforcing existing educational and socioeconomic inequalities, underscoring the need for transparency, regulation, and oversight in the use of genAI within HE decision-making.

**Highlights:**

- GenAI is widely used by HE applicants despite little study of its biases.
- 216 prompts across 3 chatbots generated 3240 programme suggestions.
- Grades and priorities drove major shifts in language and recommendations.
- Gender-coded wording mapped onto research strength and entry standards.
- GenAI biases may reinforce inequalities in HE admissions decision-making.

## 1. Introduction

Use of generative artificial intelligence (genAI) models by the public has dramatically increased over recent years, with an estimated 50% of the population currently using them (Rainie, 2025). GenAI platforms, such as ChatGPT, Gemini, and Copilot, generate information by processing text (large language models; LLM) and other media available across the internet. GenAI systems have demonstrated broad applicability across fields including education (Francis et al., 2025), medicine (Rouzrokh et al., 2025), software engineering (Rodriguez et al., 2024), and economics (Mo & Ouyang, 2025), suggesting they may help address several challenges faced by contemporary society. A recent report analysed temporal patterns of genAI use from 37.5 million conversations, identifying the topic of work and careers advice peaking between April-June for desktop users, and education ranking within the top 10 topics searched in the June-September period (Costa-Gomes et al., 2025). GenAI use is similarly widespread amongst student populations, with a recent report suggesting over half of higher education (HE) students in the UK use genAI tools, and 83% expecting this to rise (Arowosegbe et al., 2024). Similar trends have been reported in high school students, with a Norwegian study reporting 44% using genAI in 2023, rising sharply to 71% the following year (Bueie et al., 2025). Interestingly, attitudes towards the use of genAI in their studies became more negative in subsequent years, as well as being more negative in both girls and students with pre-existing positives associations with writing. Nonetheless, genAI has the potential to support students in a wide range of roles, including summarising taught content, simplifying complex topics, planning and structuring written work, providing feedback, and even evidence for supporting intellectual and social-emotional outcomes depending on the level of study (Zhu et al., 2025).

Despite potential applications throughout education, the use of genAI has been widely criticised for several reasons including risk of plagiarism and academic integrity lapses, copyright concerns for prompt outputs, and misinformation and bias through hallucinations and citation of non-existent sources (Bobula, 2024). Indeed, many reports indicate bias in genAI outputs, perhaps due to errors being propagated from the training data LLMs use to generate information (Allan et al., 2025). For example, ChatGPT has been shown to reinforce gender biases in clinical roles (e.g. more likely to describe doctors as ‘he’ and nurses as ‘she’; reviewed by Currie et al., 2025a), as well as providing significantly different medical advice for acute coronary syndrome depending on clinician gender and ethnicity (Zhang et al., 2023). GenAI bias is also borne from geographical disparities in data availability, perpetuating race, class, and overall positivity of countries, cities, and their neighbourhoods, via the so-called ‘silicon gaze’ (Kerche et al., 2026). This bias extends to image generation, where genAI systems have been shown to overrepresent light-skinned men in depictions of both medical students (Currie et al., 2024) and academics (Currie et al., 2025b), exceeding their real-world prevalence in these populations. An emerging area of genAI bias is in its application in guiding prospective undergraduate students (Shailya et al., 2025), although recent evidence has highlighted potential benefits in supporting student decision making more generally (Cabral & Pereira, 2025). A global study by Cingillioglu (2024) used a curated genAI chatbot to identify key factors that determine post-graduation matriculation decisions in 1200 participants. They identified several key topics, including location, ranking, the programme, and the university, which guide student destinations after graduation, confirming the utility of genAI in these decisions.

Despite extensive work on genAI bias in clinical, geographical, and image-generation contexts, little is known about its impact on HE decision-making at the undergraduate application level. In the UK, HE institutions provide annual programme information (e.g. entry requirements, programme length, research strengths) on their website, collated centrally through the Universities and Colleges Admissions Service (UCAS), allowing prospective students to browse, select, and apply for their top five choices for undergraduate study. Whilst some studies have provided qualitative assessments of, for example, gender bias in genAI-guided science, technology, engineering, and mathematics (STEM) careers advice (Shaner et al., 2024), a comprehensive quantitative analysis of such biases in guiding undergraduate application decision-making has not been explored. Hence, given the widespread adoption of genAI across society and within student populations, here we aimed to establish patterns of bias in guiding prospective undergraduate students.

## 2. Methods

### 2.1. Prompt design

Prompts were designed to reflect key characteristics of UK undergraduate applicants, using neuroscience as a representative degree, incorporating components that varied in applicant gender (female; male), type of student (‘study type’: current A-level student or 16-18 year old equivalent; mature student), type of education (state, private), predicted/achieved grades (BBC/112 UCAS points/2.7 GPA; AAB/136 UCAS points/3.7 GPA; A*AA/152 UCAS points/4.0 GPA), subjects studied Biology/Chemistry/Mathematics, described as ‘science heavy’; Biology/Psychology/English, described as ‘science intermediate’; and Psychology, English, Geography, described as ‘science light’), and priority (quality of neuroscience research; quality of neuroscience teaching; students’ opinions of the quality of their programme) in a fully factorial design. Each prompt included a statement asking to suggest the five most appropriate universities and a brief description explaining why. An exemplar prompt is provided: “*I want to study a BSc in Neuroscience at a UK university. I am a male mature student and studied at a state school. I have achieved A*A*A (160 UCAS points) in Biology, Chemistry, and Maths. The quality of neuroscience research is the most important factor for me when choosing where to study. Please suggest the 5 most appropriate universities I should apply to, giving a brief description of why they are suitable.*”. Hence, a total of 216 prompts were generated and each submitted to three genAI chatbots (ChatGPT-4o, Microsoft Copilot Chat, and Google Gemini Flash 2.5), resulting in 648 submitted prompts. The chatbots were intentionally selected to reflect differences in availability amongst the population being studied. Each submitted prompt/genAI pair were ordered using a random number generator, such that 58-60 were submitted each day over the course of 11 consecutive working days in July 2025, to mitigate the potential effects of day or sequence on output responses. Within each day, there was an equal distribution of all other prompt components. The 648 submitted prompts generated output responses each containing five recommendations, and therefore 3240 individual institution/programmes were generated. In cases where an individual recommendation included more than one university, prompts were immediately rerun (n=3). Full output responses were processed to remove decorative punctuation (e.g. ---, *), emojis, bullet points, and additional lines/spaces.

### 2.2 Exploratory text analysis

#### 2.2.1 Composition of output responses

Output responses were analysed for basic measures including number of words, number of characters, and mean number of characters per word.

#### 2.2.2. Readability

The overall readability of processed output responses was analysed by two measures: Flesch-Kincaid Grade Level (FKGL) and the Simple Measure of Gobbledygook (SMOG) index. FKGL uses number of words, sentences, and syllables to estimate the US reading grade required for the text (Kandula & Zeng-Treitler, 2008), whereas the SMOG index uses the number of polysyllables (words of three or more syllables) and sentences, often used to assess text for healthcare comprehension (McLaughlin, 1969):

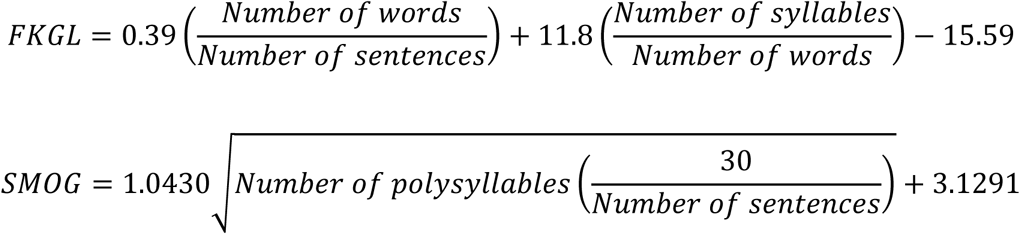

#### 2.2.3. Gender-coded language

To explore differences in gender coding of genAI output responses, a Gender Decoder tool (https://gender-decoder.katmatfield.com/) was used on full, processed output responses, based on disparities in gender-coded language in job advertisements (Gaucher et al., 2011). The Gender Decoder tool analyses the frequency of typically feminine (e.g. affectionate, collaborative, supportive) and masculine-coded (e.g. active, adventurous, aggressive) language, categorising text into ‘strongly feminine, ‘feminine’, ‘neutral’, ‘masculine’, and ‘strongly masculine’. The overall gender code for each output response was recorded.

#### 2.2.4. N-gram analysis

All output responses were aggregated and analysed for frequency of n-grams (unigrams, bigrams, trigrams). An arbitrary minimum frequency threshold of 120 was set and all resulting n-grams were screened for relevance. Conjunctions, pronouns, numbers, institution names, irrelevant bigrams/trigrams (e.g. ‘neuroscience in’, ‘would you like’), and elements of the prompt inputs (e.g. ‘neuroscience teaching’, ‘suitable’, ‘entry requirements’) were ignored. Following this, 36 of the most frequently used n-grams were identified and organised into 16 concepts across three overall categories: 1) aspect of HE (applications, research, strong neuroscience, student feedback, teaching, widening access); 2) markers of esteem (excellence, exceptional, good, quality, strengths, top, value, competitive, cutting edge, leading, outward facing, renowned, reputation, vibrant); 3) programme opportunities (clinical, interdisciplinary, opportunity, support). A full list of n-grams for each of the 16 concepts is shown in Supplementary Table 1. Frequency of n-grams within output responses was analysed with respect to gender, grades, and priority included in the prompt.

#### 2.2.5. Source analysis

Output responses were screened for commonly cited sources, where provided. The frequency of output responses that contained reference to the user-generated content (UGC) sources Reddit and Wikipedia were analysed.

### 2.2. Programme recommendations

Individual recommendations were tagged with the institution and programme. For recommended institutions without a dedicated undergraduate Neuroscience BSc programme, prompts, institution websites, and the UCAS website were consulted to identify the closest match (e.g. Biomedical Sciences BSc). Entry requirements were identified through institution websites and cross-referenced through the UCAS website, using the minimum accepted grades and excluding contextual offers.

#### 2.2.2 Markers of esteem

Institutions and their associated programmes were assessed for several markers of esteem. Institutions were tagged as to whether they were in the Russell Group (a self-selected group of 24 research-intensive public universities; described herein as ‘elite’ institutions) or Oxbridge (Universities of Oxford and Cambridge; themselves part of the Russell Group and further globally renowned for academic excellence, rigorous entrance processes, and research prestige). Markers of esteem for research (Research Excellence Framework 2021, REF21, specifically Unit of Assessment 4: Psychology, Psychiatry and Neuroscience – not compulsory and UK-wide, measured by numerical ranking/scores), teaching (Teaching Excellence Framework 2023, TEF23 – compulsory, England only, measured by award categories of bronze, silver, and gold), and student voice (National Student Survey 2024, NSS24 – compulsory and UK-wide, measured by positivity scores in %) were included to align with the applicant priorities included in prompts. NSS data was taken from July 2024, rather than July 2025, as these data were not publicly available until after starting to run the prompts. The association between gender-coded language of output responses and markers of esteem of recommended institutions was analysed.

### 2.3 Statistics

All statistics were performed using SPSS (v29.0.0.0, IBM, US) and data were visualised using GraphPad Prism (v10.2.2, US), with all bar graphs showing mean ± SEM unless otherwise stated. General linear models (GLM) were used to analyse continuous data (length of output responses, readability, REF23/NSS24 scores, grade suggestions) with relevant covariates and fixed factors used in models where appropriate. Chi-squared (χ²) tests were used to analyse categorical data (e.g. gender code, TEF23 categories) with Cramer’s V shown as a measure of effect size for statistically significant effects. For n-gram analyses, the Bonferroni correction was used to adjust the significance threshold to α=0.001 based on the number of tests (0.05/(16*3)), with significant effects denoted by ‘s’. For all other analyses, α=0.05 with significant effects denoted by ‘*’.

### 2.5 Environmental impact

To recognise the importance of sustainable and transparent use of genAI in research, the open-source Python software EcoLogits was used to assess the environmental impact of the 216 prompts submitted for this study (Rincé & Banse, 2025). EcoLogits does not include estimates for Copilot, so equivalent values for ChatGPT were used based on their similarities. The option “Write an article summary (250 output tokens)” was selected as the closest example for each of the three genAI chatbots used.

Electricity consumption, greenhouse gas emissions, energy required from primary sources, and water consumption data were scaled by the number of prompts used to generate final values. The author recognises the additional, and likely more substantial, environmental impact of genAI from model training, data storage, and predicted growth of use.

## 3 Results

### 3.1. General composition & readability

From the 3240 individual recommendations, a total of 55 institutions were specified, of which 22 (40%) were from the Russell Group, with the five most frequent accounting for 50% of all recommendations. Most institutions were based in England (n=47, 85%), with a minority in Scotland (n=6, 11%) and Wales (n=2, 4%). Only 21 (38%) had a dedicated Neuroscience BSc programme – a further 18 (33%) had Biomedical Sciences BSc as the closest programme, with the remaining 16 programmes (29%) offering combinations of psychology, natural sciences, and medical bioscience-related programmes. Entry requirements ranged from 88-160 UCAS points (equivalent to 1.7-4.0 GPA), with mean, median, and mode values of 124 (3.2 GPA), 120 (3.0 GPA), and 112 (2.7 GPA), respectively. For markers of esteem, data was available for n=49 institutions (89%) for REF21, n=47 (85%) for TEF23, and n=55 (100%) for NSS24.

The day prompts were submitted significantly affected general composition (GLM characters, F_10,637_=2.2, p=0.017; GLM words, F_10,637_=2.2, p=0.015; GLM characters/words, F_10,637_=2.2, p=0.018; Supplementary Figure 1A-C) and readability (GLM FKGL, F_10,637_=3.2, p<0.001; GLM SMOG, F_10,637_=3.0, p=0.001; Supplementary Figure 1D-E) of output responses, likely driven by an increase on day 1 and decrease on day 2. There was no effect of day on distribution of gender-coded language (Supplementary Figure 1F).

Prompts describing science heavy subjects (Biology, Chemistry, Maths) resulted in significantly fewer characters (GLM, F_2,645_=3.0, p=0.049; Figure 1A) and words (GLM, F_2,645_=3.1, p=0.046; Figure 1B) compared to science light subjects (Psychology, English, Geography). Similarly, the genAI chatbot used significantly affected the number of characters (GLM, F_2,645_=1112.2, p<0.001; Figure 1A) and words (GLM, F_2,645_=1146.0, p<0.001; Figure 1B), with Copilot producing fewer and Gemini producing more. A higher number of characters per word was associated with prompts containing higher grades (GLM, F_2,645_=8.1, p<0.001; Figure 1C) and with research priorities (GLM, F_2,645_=7.4, p<0.001; Figure 1C), as well as from ChatGPT (GLM, F_2,645_=156.8, p<0.001; Figure 1C). A higher FKGL was significantly associated with prompts describing mature students (GLM, F_1,645_=4.5, p=0.033; Figure 1D), higher grades (GLM, F_2,645_=6.9, p=0.001; Figure 1D), research priorities (GLM, F_2,645_=10.1, p<0.001; Figure 1D), and those produced by Gemini (GLM, F_2,645_=414.9, p<0.001; Figure 1D). Similarly, a higher SMOG was significantly associated with prompts describing higher grades (GLM, F_2,645_=4.3, p=0.014; Figure 1E), research priorities (GLM, F_2,645_=11.5, p<0.001; Figure 1E), and those produced by Gemini (GLM, F_2,645_=514.0, p<0.001; Figure 1E). Input gender and school type did not affect any general text or readability measures (Figure 1A-E).

**Figure 1.**
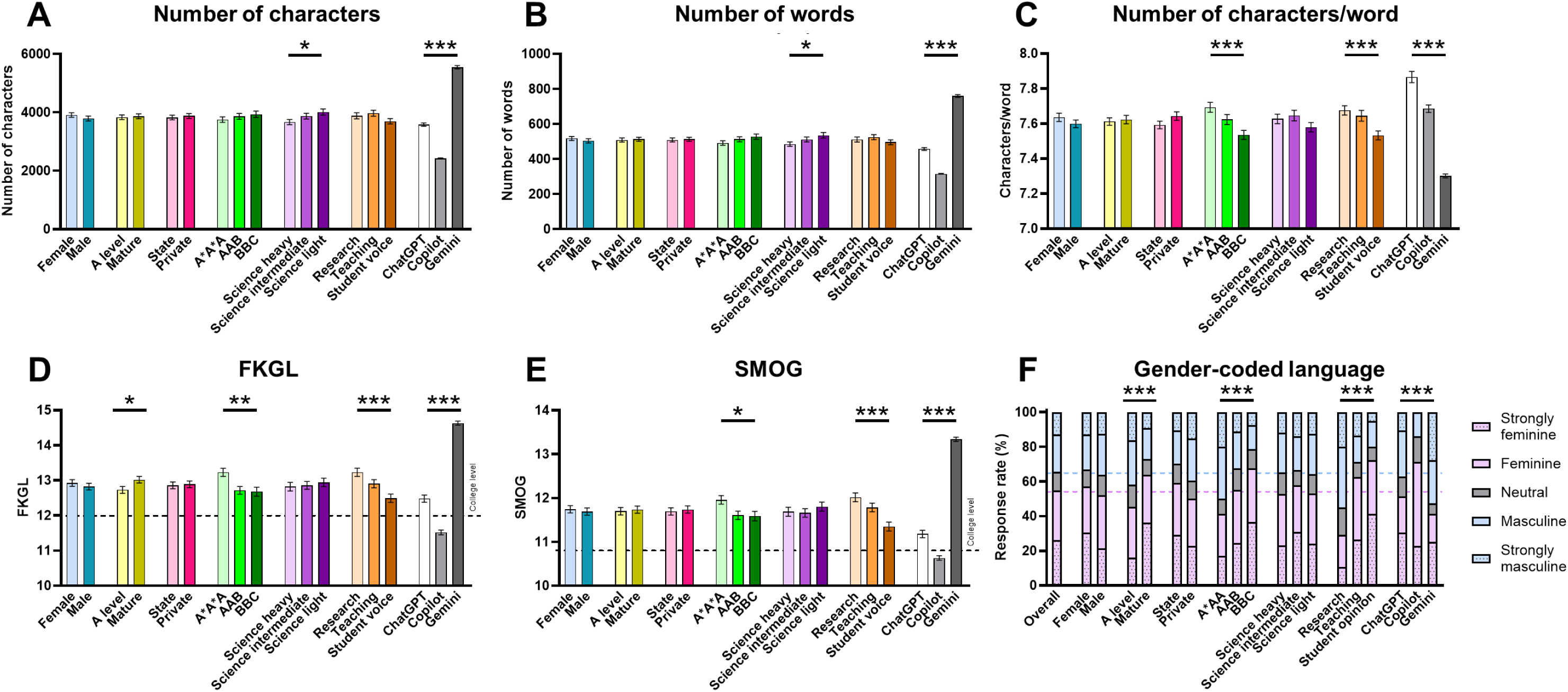
Text, readability, and gender-code analysis of output responses. The effect of prompt details (gender, study type, school type, grades, subjects, priority) and genAI used on **A)** number of characters, **B)** number of words, **C)** number of characters per word, the readability indexes **D)** FKGL and **E)** SMOG (horizontal dashed lines represent college-level reading ability), and **F)** gender-coded language (horizontal dashed lines show overall distributions of feminine and masculine-coded language across all prompts). All bars show mean ± SEM. Key: *p<0.05, **p<0.01, ***p<0.001. FKGL, Flesch-Kincaid Grade Level; SMOG, Simple Measure of Gobbledygook.

### 3.2. Gender-coded language

Overall, output responses were most likely to be feminine (n=185, 29%) or strongly feminine (n=169, 26%), followed by masculine (n=141, 22%), strongly masculine (n=83, 13%), and least likely to be neutral (n=70, 11%). The most used feminine-coded words were ‘support-’ (n=539), ‘feel-’ (n=162), ‘depend-’ (n=154), ‘collab-’ (n=132), and ‘understand-’ (n=126), whereas the most used masculine-coded words were ‘lead-’ (n=364), ‘active-’ (n=268), ‘logic-’ (n=264), ‘opinion-’ (n=114), and ‘independen-’ (n=81).

Gender-coded language of output responses was significantly affected by study type (χ²(4)=38.0, p<0.001, Cramer’s V=0.242), grades (χ²(4)=46.6, p<0.001, Cramer’s V=0.190), and applicant priority (χ²(8)=103.9, p<0.001, Cramer’s V=0.283), where A-level students, higher grades, and those with research priorities were more likely to be given masculine-coded output responses (Figure 1F). In contrast, gender-coded language was not significantly affected by input gender, school type, or subjects.

### 3.3. N-gram analysis

Overall, prompt gender significantly affected frequency of words in 1/16 concepts, grades affected that of 10/16 concepts, and priority affected that of 15/16 concepts (Figure 2A). Prompts describing male applicants were significantly more likely to produce output responses citing ‘interdisciplinary’ compared to females (GLM, F_1,646_=10.4, p=0.001; Figure 2B). No other n-grams were significantly affected by gender described in the prompt.

**Figure 2.**
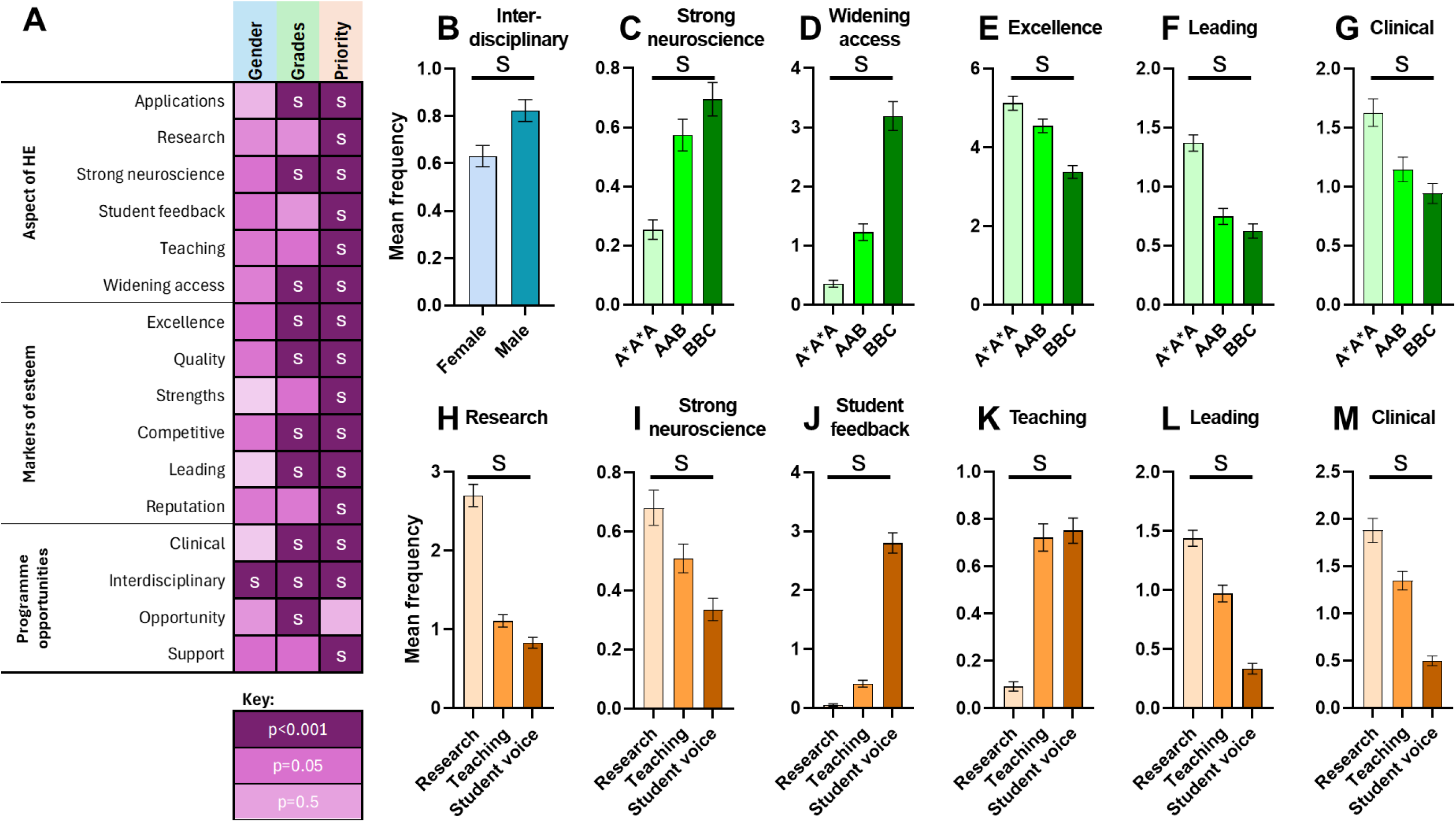
N-gram analysis of output responses. **A)** Summary of tests run, using input variable of gender (blue), grades (green), and priority (orange) across the 16 identified n-grams, grouped into three categories. Heat map (purple) shows p-values from 42 general linear models, with the Bonferroni-adjusted threshold of p<0.001 (represented by ‘s’). Dark purple with a white ‘s’ represents significant effects. Bar graphs show a selection of key findings with significant effects of prompt **B)** gender, **C-G)** grades, and **H-M)** priority. All bars show mean ± SEM. Key: ^s^p<0.001.

Several n-grams were significantly associated with prompts describing lower grades, including ‘strong neuroscience’ (GLM, F_2,645_=21.3, p<0.001; Figure 2C) and ‘widening access’ (GLM, F_2,645_=76.0, p<0.001; Figure 2D), with similar effects seen for ‘applications’ and ‘quality (Supplementary Figure 1G-H). In contrast, higher input grades were significantly associated with ‘excellence’ (GLM, F_2,645_=26.9, p<0.001; Figure 2E), ‘leading’ (GLM, F_2,645_=37.2, p<0.001; Figure 2F), and ‘clinical’ (GLM, F_2,645_=11.8, p<0.001; Figure 2G), with the same pattern seen for ‘competitive’, ‘interdisciplinary’, and ‘opportunity’ (Supplementary Figure 1I-K).

For the effect of applicant priority, those stating research quality were significantly associated with emphasis on ‘research’ (GLM, F_2,645_=100.9, p<0.001; Figure 2H), ‘strong neuroscience’ (GLM, F_2,645_=12.7, p<0.001; Figure 2I), ‘leading’ (GLM, F_2,645_=80.9, p<0.001; Figure 2L), and ‘clinical’ (GLM, F_2,645_=51.6, p<0.001; Figure 2M). In contrast, prompts stating that student voice was most important were significantly more likely to result in outputs describing ‘student feedback’, whilst this was almost absent from those stating that research or teaching was a priority (GLM, F_2,645_=198.8, p<0.001; Figure ^2^J). Output responses referring to ‘teaching’ concepts were significantly associated with prompts that specified either teaching or student voice as their priority, whilst this was almost absent in those stating research priorities (GLM, F_2,645_=62.4, p<0.001; Figure 2K). Applicant priority also had a range of significant effects on output responses referring to the concepts of ‘applications’, ‘widening access’, ‘excellence’, ‘quality’, ‘strengths’, ‘competitive’, ‘reputation’, ‘interdisciplinary’, and ‘support’ (Supplementary Figure 1L-T).

### 3.4. Reference analysis

Output responses cited a range of sources, including official institution websites (.ac.uk) and UCAS (https://www.ucas.com/), as well as additional sources summarised in Table 1.

**Table 1.** Summary of sources provided in genAI output responses.

| Title | Link provided |
| --- | --- |
| **Unifresher** | https://unifresher.co.uk/uni-prep/rankings/best-universities-for-neuroscience/ |
| **The Complete University Guide** | https://www.thecompleteuniversityguide.co.uk/courses/search/undergraduate/neuroscience |
| **Edurank** | https://edurank.org/biology/neuroscience/gb/ |
| **What Uni** | https://www.whatuni.com/degree-courses/search?subject=neuroscience |
| **Uniplus Global** | https://uniplusglobal.com/blog/504/top-neuroscience-schools-in-the-uk-2025-university-rankings-unveiled/ |
| **Go Study In** | https://gostudyin.com/study-in-uk/study-guides/top-universities-to-study-neuroscience/ |
| **University Guru** | https://www.universityguru.com/-uk/neuroscience |
| **Brittania Study** | https://britannia-study.co.uk/universities/best-universities-for-neuroscience/ |
| **Research.com** | https://research.com/university-rankings/neuroscience/gb |
| **US News** | https://www.usnews.com/education/best-global-universities/united-kingdom/neuroscience-behavior |
| **Student Crowd** | https://www.studentcrowd.com/ |

In addition to these sources, many output responses referenced UGC including Reddit and Wikipedia. Based on potential issues with reliability and accuracy of information, the frequency of these sources was analysed. Both Reddit and Wikipedia were almost exclusively cited by ChatGPT and were significantly more likely to be cited in prompts describing female applicants. Similarly, Reddit was significantly more likely to be cited in output responses to prompts describing privately educated applicants, as well as those with research or teaching priorities compared to student voice (Table 2).

**Table 2.** Effect of prompt component on frequency of user-generated content (UGC) citations in output response recommendations. Results in bold show significant findings. Key: *p<0.05, **p<0.01, ***p<0.001. V, Cramer’s V.

| Prompt component |  | Reddit |  | Wikipedia |  |
| --- | --- | --- | --- | --- | --- |
| | | # output responses citing (%) | $\chi^2$ results | # output responses citing (%) | $\chi^2$ results |
| Gender | Female | **44**<br>**(71%)** | **$\chi^2(1)=12.1$**<br>**\*\*\*p<0.001**<br>**V=0.136** | **53**<br>**(77%)** | **$\chi^2(1)=22.2$**<br>**\*\*\*p<0.001**<br>**V=0.185** |
|  | Male | **18**<br>**(29%)** |  | **16**<br>**(23%)** |  |
| Study type | A level | 31<br>(50%) | $\chi^2(1)=0.0$<br>p=1.000 | 30<br>(43%) | $\chi^2(1)=1.3$<br>p=0.252 |
|  | Mature | 31<br>(50%) |  | 39<br>(57%) |  |
| School | State | **21**<br>**(34%)** | **$\chi^2(1)=7.1$**<br>**\*\*p=0.008**<br>**V=0.105** | 29<br>(42%) | $\chi^2(1)=2.0$<br>p=0.161 |
|  | Private | **41**<br>**(66%)** |  | 40<br>(58%) |  |
| Grades | A*A*A | 26<br>(42%) | $\chi^2(2)=3.2$<br>p=0.197 | 28<br>(41%) | $\chi^2(2)=3.8$<br>p=0.150 |
|  | AAB | 21<br>(34%) |  | 25<br>(36%) |  |
|  | BBC | 15<br>(24%) |  | 16 (23%) |  |
| Subjects | Science heavy | 20<br>(32%) | $\chi^2(2)=0.5$<br>p=0.793 | 24<br>(35%) | $\chi^2(2)=0.3$<br>p=0.864 |
|  | Science intermediate | 23<br>(37%) |  | 21<br>(30%) |  |
|  | Science light | 19<br>(31%) |  | 24<br>(35%) |  |
| Priority | Research | **23**<br>**(37%)** | **$\chi^2(2)=8.2$**<br>**\*p=0.017**<br>**V=0.112** | 25<br>(36%) | $\chi^2(2)=0.4$<br>p=0.823 |
|  | Teaching | **28**<br>**(45%)** |  | 23<br>(33%) |  |
|  | Student voice | **11**<br>**(18%)** |  | 21<br>(30%) |  |
| GenAI | ChatGPT | **58**<br>**(94%)** | **$\chi^2(2)=112.3$**<br>**\*\*\*p<0.001**<br>**V=0.416** | **69**<br>**(100%)** | **$\chi^2(2)=154.4$**<br>**\*\*\*p<0.001**<br>**V=0.488** |
|  | Copilot | **0**<br>**(0%)** |  | **0**<br>**(0%)** |  |
|  | Gemini | **4**<br>**(6%)** |  | **0**<br>**(0%)** |  |

### 3.5. Markers of esteem

#### 3.5.1. Institution recommendation

Overall, Russell group institutions were significantly more likely to be recommended for prompts that described mature (χ²(2)=14.4, p<0.001, V=0.067), privately educated students (χ²(2)=9.6, p=0.008, V=0.054) with high grades (χ²(3)=498.5, p<0.001, V=0.392), studying science subjects (χ²(3)=9.6, p=0.022, V=0.054), and with research priorities (χ²(3)=111.2, p<0.001, V=0.185; Figure 3A, effects shown by ‘*’). Similarly, output responses were significantly more likely to recommend Oxbridge to female (χ²(2)=6.0, p=0.049, V=0.043), mature students (χ²(2)=6.6, p=0.037, V=0.045), with higher grades (χ²(3)=225.6, p<0.001, V=0.264) and research priorities (χ²(3)=151.3, p<0.001, V=0.216; Figure 3A, effects shown by ‘#’).

**Figure 3.**
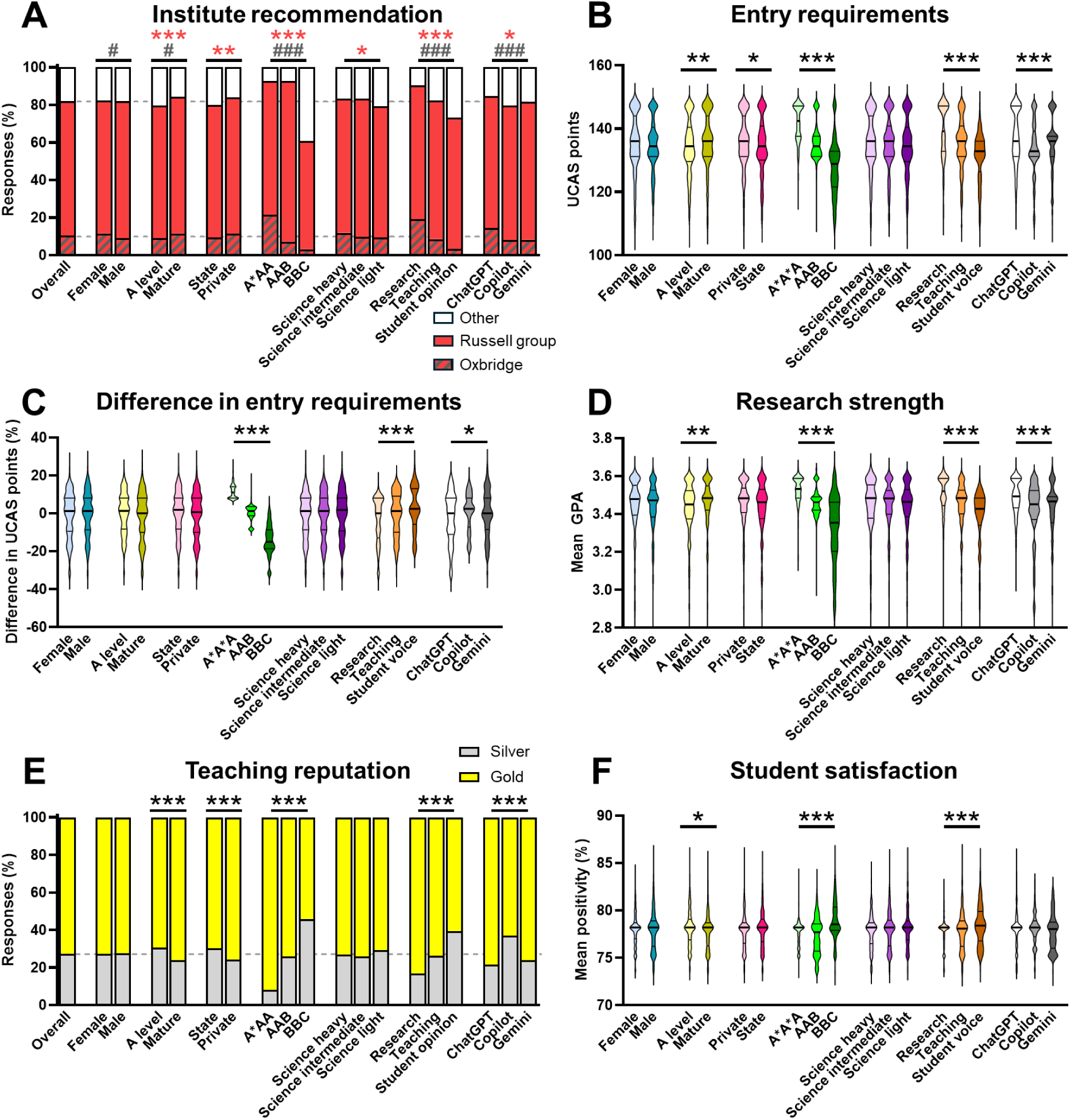
Effect of prompt components on markers of esteem of output recommendation. **A)** Institution recommendations within the Russell (red) and Oxbridge (red hashed) groups, compared to other institutions (white; horizontal grey dashed lines show values for all output responses combined). Effect of prompt components on **B)** entry requirements of recommended institutions; **C)** difference in entry requirements of recommended institutions compared to input grades; **D)** research strength (as measured by the REF21 GPA for neuroscience); **E)** teaching reputation (as measured by TEF23 for student outcomes; horizontal grey dashed lines show values for all output responses combined); and **F)** student satisfaction (as measured by NSS24 overall positivity). For A, red ‘*’ symbols represent significant effects on Russell group recommendations, whereas grey ‘^#^’ symbols represent significant effects on Oxbridge recommendations (^#^p<0.05, ^###^p<0.001). For all others, *p<0.05, **p<0.01, ***p<0.001. For B-D and F, violin plots show median and quartiles. Legends shown in panels A and E.

#### 3.5.2. Entry requirements

The mean entry requirements of recommended institutions were significantly higher for mature (GLM, F_1,646_=7.8, p=0.005), privately educated students (GLM, F_1,646_=4.2, p=0.041) with higher grades (GLM, F_2,645_=197.8, p<0.001) and research priorities (GLM, F_2,645_=50.9, p<0.001; Figure 3B). When analysing the mean difference in entry requirements (that is, the difference in entry requirements from the recommended institutions compared to the prompt grades), a similar pattern emerged. Entry requirements were significantly inflated compared to prompts for those with higher grades (GLM, F_2,645_=1093.0, p<0.001) and, interestingly, underinflated for those with research priorities (GLM, F_2,645_=14.4, p<0.001; Figure 3C). Input gender and subjects had no effect on entry requirements of difference in entry requirements of recommendations.

#### 3.5.3. Research strength

Prompts describing mature students (GLM, F_1,646_=8.0, p=0.005), higher grades (GLM, F_2,645_=148.4, p<0.001), and research priorities (GLM, F_2,645_=36.1, p<0.001) were significantly associated with increased research strength as measured by the REF21 GPA for neuroscience (Figure 3D). Similar patterns were seen for a secondary measure of research strength (REF21 rank for neuroscience) where study type (GLM, F_1,646_=7.7, p=0.006), grades (GLM, F_2,645_=159.3, p<0.001), and priority (GLM, F_2,38.9_, p<0.001) all had significant effects (Supplementary Figure 2A).

#### 3.5.4. Teaching reputation

Recommended institutions were significantly more likely to have higher teaching prestige (as indicated by a higher proportion of ‘gold’ for student outcomes in TEF23) for prompts describing mature (χ²(2)=16.8, p<0.001, V=0.081), privately educated students (χ²(2)=14.4, p<0.001, V=0.075), with higher grades (χ²(3)=312.7, p<0.001, V=0.351) and prioritising the student voice (χ²(3)=109.7, p<0.001, V=0.208; Figure 3E). Similar significant effects were found for other aspects of teaching excellence (as measured by TEF23), including student experience where study type (χ²(4)=10.0, p=0.041, V=0.063), grades (χ²(6)=246.3, p<0.001, V=0.220), and priority (χ²(6)=25.5, p<0.001, V=0.071) had significant effects (Supplementary Figure 2B), and the overall score where input grades (χ²(3)=37.3, p<0.001, V=0.121) and priority (χ²(3)=23.0, p<0.001, V=0.095) had significant effects (Supplementary Figure 2C). Gender and subjects had no effect on markers of teaching reputation.

#### 3.5.5. Student satisfaction

Prompts describing A-level students (GLM, F_1,646_=5.8, p=0.016), lower grades (GLM, F_2,645_=46.8, p<0.001), and those prioritising student voice (GLM, F_2,645_=9.0, p<0.001) were associated with significantly increased student satisfaction (as measured by NSS24 positivity; Figure 3F). However, importantly the differences in means were small, at 0.4% for study type (A-level vs mature students), 1.6% for grades (BBC vs AAB), and 0.8% for priority (student voice vs research).

#### 3.5.6. Effect of chatbot

Finally, the chatbot used also significantly affected recommendations of both Russell group (χ²(3)=10.5, p=0.014, V=0.057; Figure 3A) and Oxbridge (χ²(3)=32.1, p<0.001, V=0.099), as well as the mean entry requirements (GLM, F_2,645_=11.5, p<0.001; Figure 3B), difference in entry requirements (GLM, F_2,645_=4.5, p=0.012; Figure 3C), research strengths (GLM, F_2,645_=18.8, p<0.001; Figure 3D), and teaching reputation (χ²(3)=20.4, p<0.001, V=0.090; Figure 3E) of the recommended institutions, without affecting that of student satisfaction scores (Figure 3F).

### 3.6. Association between gender-coded language on markers of esteem

Feminine-coded language was significantly associated with reduced entry requirements of the recommended institutions in output responses (GLM, F_4,643_=26.5, p<0.001; Figure 4A). There was no overall effect of gender coding on mean difference in entry requirements (GLM, F_4,643_=2.3, p=0.054; Figure 4B, left-hand side). However, when categories were collapsed into overall feminine (strongly feminine + feminine) and overall masculine (strongly masculine + masculine), there was a significant association such that genAI output responses were more likely to provide masculine-coded recommendations when entry requirements exceeded the applicants grades, with the reverse true for feminine and neutral-coded output responses (GLM, F_2,645_=3.6, p=0.028; Figure 4B, right-hand side). Similar patterns were observed with institution markers of research esteem, where masculine-coded recommendations were significantly associated with institutions with stronger neuroscience research, including research GPA (GLM, F_4,643_=19.1, p<0.001; Figure 4C) and rank (GLM, F_4,643_=20.3, p<0.001; Figure 4D). In contrast, genAI output responses that were feminine-coded were significantly more likely to recommend institutions with better student satisfaction scores (GLM, F_4,643_=6.4, p<0.001; Figure 4E).

**Figure 4.**
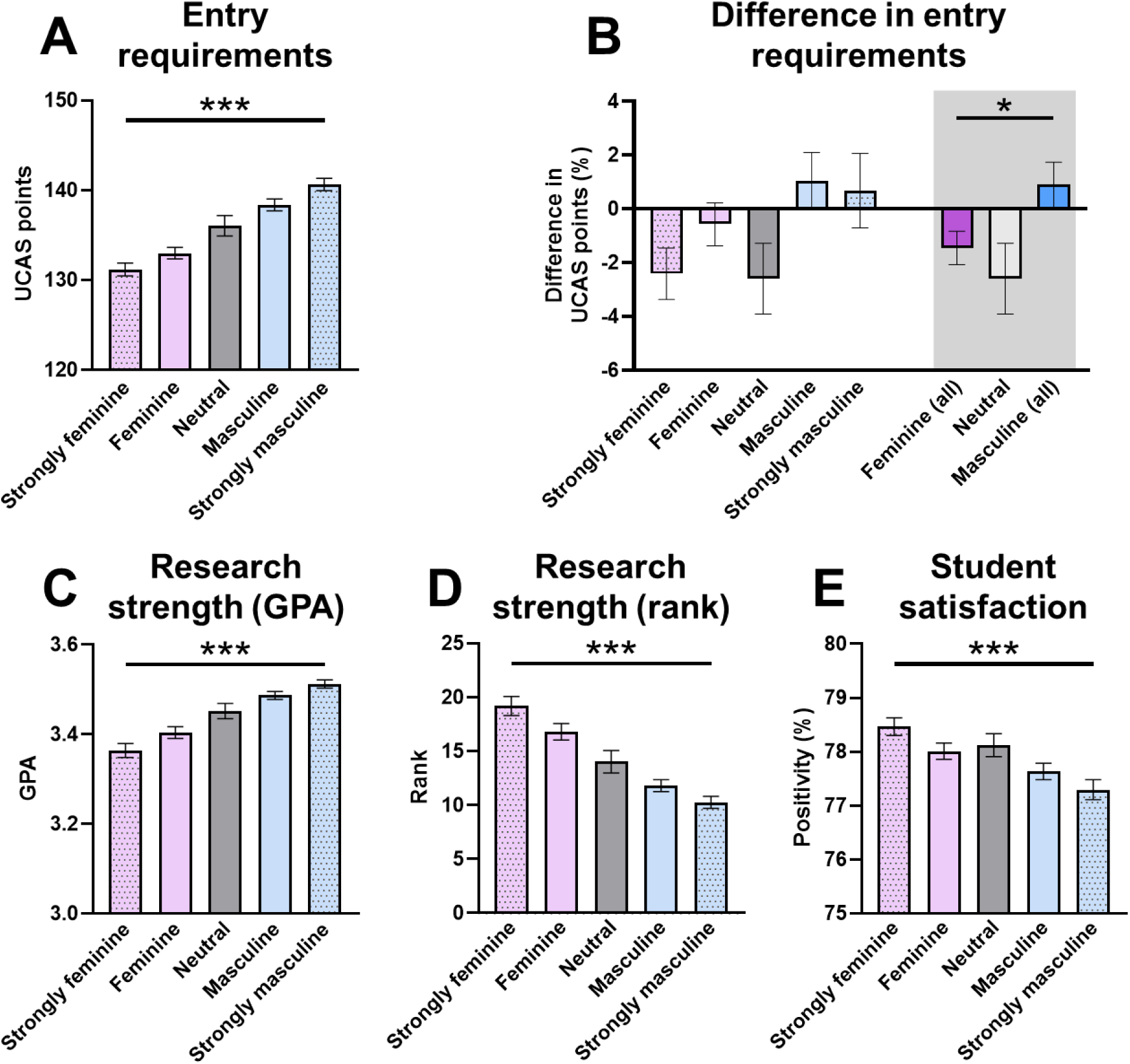
Association of gender-coded language and markers of esteem in output responses. Effect of gender-coded language in output responses on markers of esteem averaged across the five recommended institutions. **A)** Entry requirements for recommended programmes, measured by UCAS points. **B)** Difference in entry requirements in recommended programmes compared to applicant input grades (note the two grey bars for ‘Neutral’ show the same data). Research strength of recommended institutions, measured by the REF21 neuroscience research **C)** GPA and **D)** rank. **E)** Student satisfaction of recommended programmes based on overall NSS24 positivity scores. All bars show mean ± SEM. Key: *p<0.05, ***p<0.001. GPA, grade point average; UCAS, University and Colleges Admissions Service.

### 3.7. Environmental impact statement

In total, the prompts used in this study are estimated to have used *either*: a) 29.2 Wh of electricity (equivalent to ∼30 min use of a standard incandescent lightbulb); b) 11.6 gCO_2_eq greenhouse gas emissions (equivalent of a standard unleaded car driving ∼71 m); c) 287 kJ of primary energy (equivalent to the energy content of an egg); *or*, d) 126 mL water (equivalent to ∼1/2 a cup).

## 4 Discussion

Here, we investigated patterns of bias in genAI outputs in guiding prospective undergraduate students in decision making for HE studies. We uncover several critical factors that guide genAI outputs in this context, with particularly strong effects of applicant grades and priorities when choosing where to study, which skewed the recommended institutions with relation to markers of esteem. These factors were also associated with significant changes in the prevalence of gender-coded language, independently of the applicant gender, which further skewed markers of esteem of recommended institutions.

### 4.1. Applicant grades and priorities affect gender-coded language and contents of genAI outputs

We first demonstrate that the grades of a prospective applicant (either high, medium, or low), as well as their stated priorities when choosing a programme to study on (research-intensive, teaching focussed, or positive student feedback) are of particular importance in determining the content of genAI outputs. While there were subtle but significant effects of these input prompt characteristics on readability (i.e. FKGL, SMOG), perhaps of more interest are the effects on gender-coded language. We show that prompts are significantly more likely to be masculine-coded for applicants who are current A-level students, those with higher grades, and with more research-focussed priorities. LLMs are known to be systematically influenced by both intrinsic biases and external social stereotypes, suggesting that while input semantics can skew outputs, wider biases may still be propagated from training data (Liu & Li, 2024) through to the response provided to users (Wu et al., 2025). This is further supported by Gaebler et al. (2025), who show that despite constant input qualifications, LLMs reinforce gender and race biases during the hiring process for teachers, with models favouring white women. This may explain why gender had minimal significant effects across our study, despite seeing effects on gender-coded outputs. Interestingly, our data demonstrated that most output responses were feminine-coded (55% compared to 35% masculine-coded), whilst wider literature suggests that LLMs default towards a masculine bias in English (Bartl et al., 2025) which can be stronger in other languages (e.g. German). Nonetheless, this may help to explain why higher-grade applicants and those with priorities in research prestige were significantly more likely to produce masculine-coded outputs.

### 4.2. Composition of prompts impacts quality of undergraduate study recommendations

We next report a range of effects of how prospective applicant characteristics affect the quality of study recommendations provided, focussing on a selection of markers of esteem across the HE sector. For example, male applicants were more likely to have the interdisciplinary nature of programmes emphasised in their recommendations. Perhaps encouragingly, applicants with higher grades had institution excellence emphasised, whereas those with lower grades were directed to widening participation information, and applicant priorities were largely reflected accurately in prompt outputs. Similarly, whilst expected effects were seen whereby applicants with higher grades and research priorities were more likely to be recommended elite, research-intensive institutions, this was also seen for those from privately educated backgrounds. Geographic, demographic, and socioeconomic biases have been demonstrated elsewhere (Shailya et al., 2025), raising concern about the long-term effects on socioeconomic mobility. Such bias in LLMs, although perhaps subtle, are likely to accumulate and perpetuate long-term structural inequalities across graduate populations. Interestingly, Shailya et al. (2025) also demonstrate that institutions in the Global North are significantly more frequently recommended by LLMs, emphasising that the structural inequalities and biases we report here are also more widely seen outside the limits of the UK. Repeating our study in the context of international study would widen the interpretation of these biases, which is of critical importance for a sector like HE which relies on transnational movement of knowledge and students.

### 4.3. GenAI perpetuates gender-coded biases in academic markers of esteem

Finally, we demonstrate that the gender-coded language of genAI output responses (which itself is skewed by input applicant characteristics) is strongly associated with academic markers of esteem. Masculine-coded outputs were significantly associated with higher programme entry requirements, a higher difference in entry requirements (i.e. more likely to suggest a programme which exceeded the applicant’s attainment), and recommendations of institutions with higher research prestige but lower student satisfaction. Soundararajan & Delany (2024) have similarly demonstrated that specific chatbots (e.g. ChatGPT) exhibit clear gender biases, reinforcing stereotypes by more frequently associating masculine-coded terms with increased competence, agency, and prestige. In contrast, our finding that feminine-coded responses are more likely to provide recommendations where student feedback is more positive may be explained by the psycholinguistic observation that females value communal and social aspects more highly (Hsu et al., 2021). Such gender biases exist even across more ‘modern’ LLMs (e.g. Llama 3, Gemma), producing gender-skewed lexical and thematic content depending on the context (Rickman, 2025). This suggests the findings we report here are not isolated to the HE sector, but span across many domains of society. Furthermore, social bias and fairness amongst LLMs are ongoing problems, with risk of ‘bias drift’, given that these phenomena vary depending on training corpora and model versions (Gallegos et al., 2024).

## 5 Conclusion

GenAI is rapidly developing as a useful tool for streamlining daily tasks and summarising large quantities of data that drive decision making, including in prospective undergraduate students. We show here that applicant characteristics, which are likely used to form prompts that guide these decisions, generate clear patterns of bias in output responses. This in turn is likely to perpetuate existing biases, with respect to gender, educational background, and socioeconomic status, working antagonistically with efforts to promote widening participation in HE. These biases may extend more widely on the international scale, such as by disproportionately advantaging HE institutions in the Global North, which may impede social mobility and internationalisation in HE. The current study therefore has widespread implications for policymakers in HE, who must respond to changing trends in how prospective undergraduate applicants review the available information on programmes and their institutions. Given such chatbots will likely remain a commonly used tool amongst prospective students for assessing potential study options, both applicants and HE institutions remain important stakeholders in the growing need for regulatory oversight around fairness, privacy, and transparency of genAI in education.

## Acknowledgements

Thanks to Dr Beatriz Costa Gomes for her thoughtful guidance and insights on the manuscript.

## Declaration

The author has no competing interests to declare.

## Supplementary Information

**Supplementary Table 1.** Concepts and n-grams used for text frequency analysis.

| Category | Concept | N-grams |
| --- | --- | --- |
| Aspect of HE | Applications | personal statement |
|  | Research | research excellence, research output, research strength, strong research, research intensive |
|  | Strong neuroscience | strong neuroscience |
|  | Student feedback | national student survey, NSS, student experience |
|  | Teaching | excellent teaching, learning environment |
|  | Widening access | accessible, contextual |
| Markers of esteem | Excellence | excellence, excellent, exceptional, top |
|  | Quality | quality, value, good, vibrant |
|  | Strengths | strong, strengths |
|  | Competitive | competitive |
|  | Leading | cutting edge, cutting-edge, leader |
|  | Reputation | globally, world class, world-class, highly regarded, renowned, reputation |
| Programme opportunities | Clinical | clinical |
|  | Interdisciplinary | interdisciplinary |
|  | Opportunity | opportunities |
|  | Support | support |

**Supplementary Figure 1.**
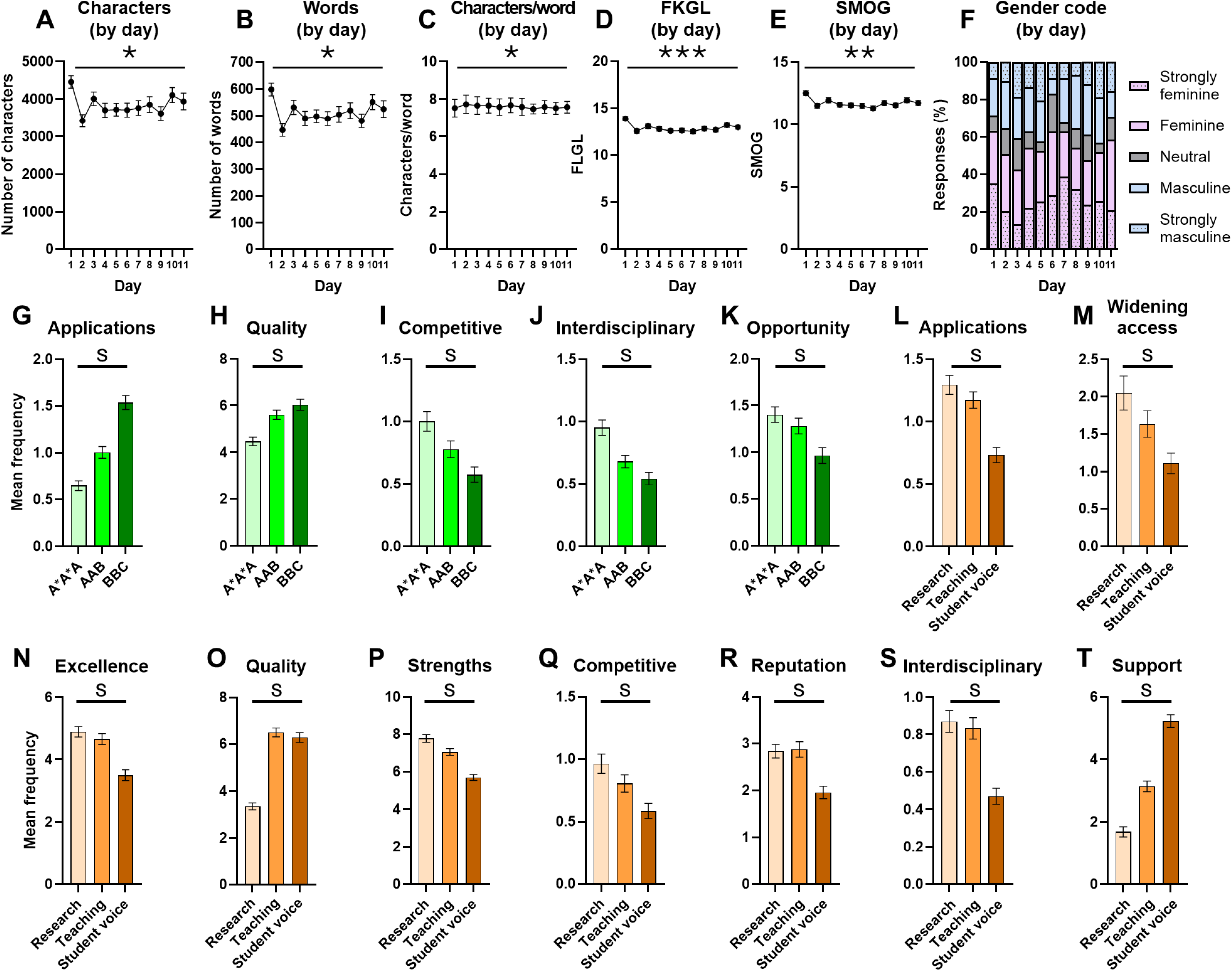
Effect of day on output response and additional n-gram analyses. Effect of the day each prompt was run on **A)** number of characters, **B)** number of words, **C)** number of words per character, readability indexes **D)** FKGL and **E)** SMOG, and **F)** gender-coded language in output responses. Effect of input prompt grades on output responses containing reference to the concepts of **G)** ‘applications’, **H)** ‘quality’, **I)** ‘competitive’, **J)** ‘interdisciplinary’, and **K)** ‘opportunity. Effect of prompt priority on output responses containing reference to the concepts of **L)** ‘opportunity’, **M)** ‘widening access’, **N)** ‘excellence’, **O)** ‘quality’, **P)** ‘strengths’, **Q)** ‘competitive’, **R)** ‘reputation’, **S)** ‘interdisciplinary’, and **T)** ‘support’. For A-F, data points show day means; for G-T, bars shown group means; for G-T, statistical tests are subject to a Bonferroni-adjusted α threshold, with significant findings indicated by ‘s’; for all, error bars show ± SEM. Key: *p<0.05, **p<0.01, ***p<0.001, ^s^p<0.001 (Bonferroni-adjusted).

**Supplementary Figure 2.**
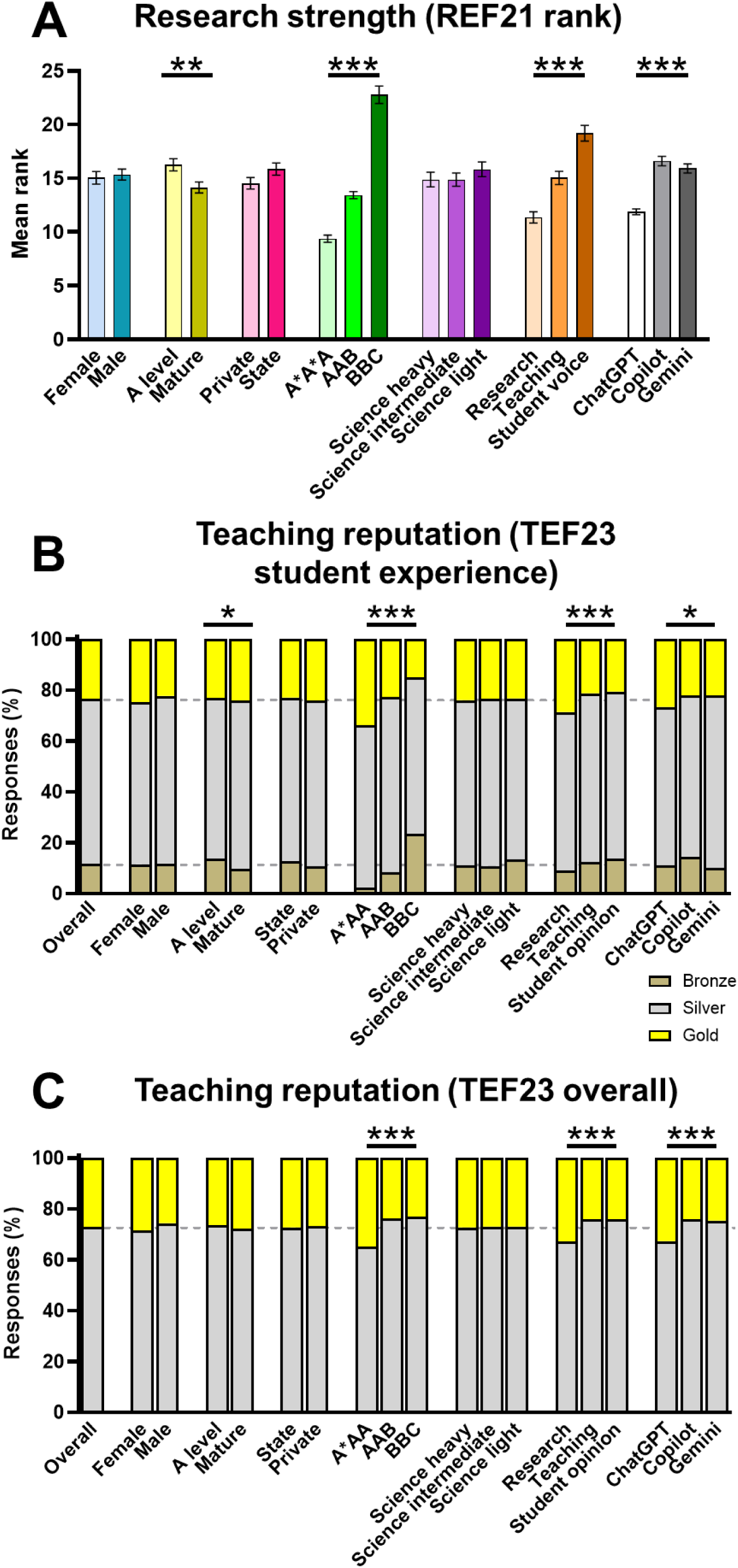
Effect of prompt components on additional markers of esteem. Effect of prompt components on **A)** research strength (as measured by REF21 rank for neuroscience research; note increased data points represent a lower rank) and teaching reputation, as measured by TEF23 for **B)** student experience and **C)** overall (for B and C, horizontal grey dashed lines show values for all output responses combined). For A, bars show mean ± SEM. Key: *p<0.05, **p<0.01, ***p<0.001. REF, Research Excellence Framework; TEF, Teaching Excellence Framework.

## References

Allan, K., Azcona, J., Sripada, S., Leontidis, G., Sutherland, C. A. M., Phillips, L. H., Martin, D. (2025). Stereotypical bias amplification and reversal in an experimental model of human interaction with generative artificial intelligence. Royal Society Open Science, 12:241472.

Arowosegbe, A., Alqahtani, J. S., Oyelade, T. (2024). Perception of generative AI use in UK higher education. Frontiers in Education, 9.

Bartl, M., Murphy, T. B., Leavy, S. (2025). Adapting Psycholinguistic Research for LLMs: Gender-inclusive Language in a Coreference Context. arXiv, 2502.13120v1.

Bobula, M. (2024). Generative artificial intelligence (AI) in higher education: a comprehensive review of challenges, opportunities, and implications. Journal of Learning Development in Higher Education, 30:112–127.

Bueie, A. A., Skar, G. B., Graham, S. (2025). High school students’ use and beliefs about generative artificial intelligence and writing in school. Reading and Writing, 10.1007/s11145-025-10702-3.

Cabral, C. A. A., Pereira, R. (2025). Digital Enhancement: Applying Generative AI in Students Decision Making Process. Digital Technologies and Transformation in Business, Industry and Organizations, 123-141.

Cingillioglu, I. (2024). What impacts matriculation decisions? Identifying students’ university choice factors on a global scale with Artificial Intelligence. Studies in Higher Education, 49:2637–2655.

Costa-Gomes, B., Chen, S., Hsueh, C., Morgan, D., Schoenegger, P., Shah, Y., Way, S., Zhu, Y., Adeline, T., Bhaskar, M., Suleyman, M., Spielman, S. (2025). It’s About Time: The Copilot Usage Report 2025 The Temporal and Modal Dynamics of Copilot Usage. Arxiv, 2512.11879v1.

Currie, G. M., Hawk, K. E., Rohren, E. M. (2025a). Generative Artificial Intelligence Biases, Limitations and Risks in Nuclear Medicine: An Argument for Appropriate Use Framework and Recommendations. Seminars in Nuclear Medicine, 55:423–436.

Currie, G. M., Hewis, J., Wheat, J. (2025b). Gender and ethnicity representation of university academics by generative artificial intelligence using DALL-E 3. Journal of Further and Higher Education, 49:1064–1078.

Francis, N. J., Jones, S., Smith, D. P. (2025). Generative AI in Higher Education: Balancing Innovation and Integrity. British Journal of Biomedical Science, 81:14048.

Gaebler, J. D., Goel, S., Huq, A., Tambe, P. (2025). Auditing large language models for race & gender disparities: Implications for artificial intelligence-based hiring. Behavioural Science & Policy, 11:1–10.

Gallegos, I. O., Rossi, R. A., Barrow, J., Tanjim, M. M., Kim, S., Dernoncourt, F., Yu, T., Zhang, R., Ahmed, N. K. (2024). Bias and Fairness in Large Language Models: A Survey. Computational Linguistics, 50:1097–1179.

Gaucher D, Friesen J, Kay AC (2011). Evidence that gendered wording in job advertisements exists and sustains gender inequality. Journal of Personality and Social Psychology, 101:109–128.

Hsu, N., Badura, K. L., Newman, D. A., Speech, M. E. P. (2021). Gender, “masculinity,” and “femininity”: A meta-analytic review of gender differences in agency and communion. Psychological Bulletin, 147:987–1011.

Kandula, S., Zeng-Treitlet, Q. (2008). Creating a Gold Standard for the Readability Measurement of Health Texts. AMIA Annual Symposium Proceedings, 353-357.

Kerche, F. W., Zook, M., Graham, M. (2026). The silicon gaze: A typology of biases and inequality in LLMs through the lens of place. Platforms & Society, 3.

Liu, W., Li, M. (2024). The analysis of technological ethical issues in generative artificial intelligence. J. Artif. Intell. Pract. 7:155–160.

McLaughlin, G. H. (1969). SMOG Grading-a New Readability Formula. Journal of Reading, 12:639–646.

Mo, H., Ouyang, S. (2025). (Generative) AI in Financial Economics. Journal of Chinese Economic and Business Studies, 23:509–587.

Rainie, L. (2025). Close encounters of the AI kind: The increasingly human-like way people are engaging with language models. Imagining the Digital Future Center. Available at: https://imaginingthedigitalfuture.org/wp-content/uploads/2025/03/ITDF-LLM-User-Report-3-12-25.pdf (accessed 28th January 2026).

Rickman, S. (2025). Evaluating gender bias in large language models in long-term care. BMC Medical Informatics and Decision Making, 25:274.

Rincé S, Banse A (2025). EcoLogits: Evaluating the Environmental Impacts of Generative AI. Journal of Open Source Software, 10:7471.

Rodriguez, D. V., Lawrence, K., Gonzalez, J., Brandfield-Harvey, B., Xu, L., Tasneem, S., Levine, D. L., Mann, D. (2024). Leveraging Generative AI Tools to Support the Development of Digital Solutions in Health Care Research: Case Study. JMIR Human Factors, 11:e52885.

Rouzrokh, P., Khosravi, B., Faghani, S., Moassefi, M., Shariatnia, M. M., Rouzrokh, P., Erickson, B. (2025). A Current Review of Generative AI in Medicine: Core Concepts, Applications, and Current Limitations. Current Reviews in Musculoskeletal Medicine, 18:246–266.

Shailya, K., Mishra, A. K., Krishnan, G. S., Ravindran, B. (2025). Where Should I Study? Biased Language Models Decide! Evaluating Fairness in LMs for Academic Recommendations. arXiv, 2509.04498v2.

Shaner, C., Griffith, H., Rathore, H. (2024). Assessing Implicit Gender Bias in Large Language Models for STEM Career Guidance Applications. 2024 IEEE Integrated STEM Education Conference (ISEC), Princeton, NJ, USA, 2024, pp. 1-6.

Wu, W., Zhao, Q., Chen, H., Zhou, L., Lian, D., Xie, H. (2025). Exploring the Choice Behavior of Large Language Models. Findings of the Association for Computational Linguistics: ACL 2025. 5194–5214.

Zhang, A., Yuksekgonul, M., Guild, J., Zou, J., Wu, J. C. (2023). ChatGPT Exhibits Gender and Racial Biases in Acute Coronary Syndrome Management. MedRxiv, doi: 10.1101/2023.11.14.23298525

Zhu, Y., Liu, Q., Zhao, L. (2025). Exploring the impact of generative artificial intelligence on students’ learning outcomes: a meta-analysis. Education and Information Technologies, 30:16211–16239.

